# Structural rather than catalytic role for mitochondrial respiratory chain supercomplexes

**DOI:** 10.1101/2023.04.19.537447

**Authors:** Michele Brischigliaro, Alfredo Cabrera-Orefice, Susanne Arnold, Carlo Viscomi, Massimo Zeviani, Erika Fernández-Vizarra

## Abstract

Mammalian mitochondrial respiratory chain (MRC) complexes are able to associate into quaternary structures named supercomplexes (SCs), which normally coexist with non bound individual complexes. The functional significance of SCs has not been fully clarified and the debate has been centered on whether or not they confer catalytic advantages to the non-bound individual complexes. Mitochondrial respiratory chain organization does not seem to be conserved in all organisms. In fact, and differently from mammalian species, mitochondria from *Drosophila melanogaster* tissues are characterized by low amounts of SCs, despite the high metabolic demands and MRC activity shown by these mitochondria. Here, we show that attenuating the biogenesis of individual respiratory chain complexes was accompanied by increased formation of stable SCs, which are missing in *Drosophila melanogaster* in physiological conditions. This phenomenon was not accompanied by an increase in mitochondrial respiratory activity. Therefore, we conclude that SC formation is necessary to stabilize the complexes in suboptimal biogenetic conditions, but not for the enhancement of respiratory chain catalysis.

## INTRODUCTION

Mitochondria are the organelles providing most of the cellular energy in form of adenosine triphosphate (ATP) in aerobic eukaryotes. The molecular machinery responsible for energy transformation is the oxidative phosphorylation (OXPHOS) system, which is canonically composed of five multiprotein complexes embedded in the inner mitochondrial membrane. OXPHOS consists of two tightly regulated processes: electron transport and ATP synthesis. Electron transport takes place between complexes I-IV and two mobile electron carriers (coenzyme Q and cytochrome *c*). During electron transport, complexes I, III and IV pump protons from the mitochondrial matrix to the intermembrane space, generating a proton gradient that provides the protonmotive force exploited by complex V to synthesize ATP. In mammalian mitochondria, mitochondrial respiratory chain (MRC) complexes I, III and IV can interact with each other forming supramolecular structures described generally by the term ‘supercomplexes’ [1, 2]. MRC supercomplexes (SCs) can have different stoichiometries and compositions, ranging from the binding of only two complexes, such as the I_1_III_2_ and III_2_IV_1_ SCs [3, 4], to higher-order associations between complexes I, III and IV, with the SC of I_1_III_2_IV_1_ stoichiometry known as the ‘respirasome’ [1, 5–9]. Now that the association of individual MRC complexes into supramolecular structures is well established, with structures of several SC species being resolved, the debate is centered on what the functional significance of these structures might be.

Several possible roles have been proposed for SCs. First, it was suggested that the association between complex I (CI) and the obligate dimer of complex III (CIII_2_) would allow substrate channeling, sequestering a dedicated coenzyme Q (CoQ) pool and allowing a more efficient electron transfer between the two complexes, while separating this electronic route from those of FADH_2_-linked dehydrogenases (e.g. complex II) to the CIII_2_ not bound to CI [1, 10–15]. This increased efficiency would in turn decrease electron leak and, as a consequence, produce less reactive oxygen species (ROS) than the individual free complexes [16, 17]. However, the available high-resolution respirasome structures show that the distance between the CoQ binding sites in CI and CIII_2_ are far apart and exposed to the membrane, thus not supporting the notion of substrate channeling within the SC structure [4, 18]. In addition, exogenously added CoQ was necessary to sustain CI activity in the purified I_1_III_2_ SC [3], arguing against a tightly bound and segregated CoQ pool as a functional component of the SC. A large body of work from the late 1960’s to the 1980’s, resulted in the ‘random collision model’ to explain electron transfer between the respiratory chain complexes, and in the evidence that CoQ is present as an undifferentiated pool[19–23]. More recently, additional proof of the non compartmentalized electronic routes from CI to CIII_2_ and from complex II (CII) to CIII_2_, came from kinetic measurements in sub-mitochondrial particles. In these systems, MRC organization in SCs was conserved but *b*- and *c*-type cytochromes in CIII_2_ were equally accessible to CI-linked and CII-linked substrates [24], and CoQ reduced by CI in the respirasomes was able to reach and readily reduce external enzymes to the SCs [25]. In addition, growing evidence supports the notion that different MRC organizations exist *in vivo*, where varying proportions of SC vs. free complexes do not result in separate and distinct CI-linked and CII-linked respiratory activities [26–29]. This is in contrast with the idea that segregation into different types of SCs and in individual complexes is necessary for the functional interplay of the MRC, leading to the adaptation of the respiratory activity to different metabolic settings [13]. The physical proximity of CIII_2_ and CIV has also been suggested to promote faster electron transfer kinetics via cytochrome *c* [4, 30, 31], although this is a matter of debate as well [32, 33].

The second main explanation to justify the existence of SCs is that they play a structural function, stabilizing the individual complexes [34, 35] and/or serving as a platform for the efficient assembly of the complexes, with a special relevance for the biogenesis of mammalian CI [36–38].

Notably, the MRC structural organization, especially the stoichiometry, arrangements and stability of the SCs, may not be conserved in all eukaryotic species [39–42]. This is the case even within mammals, as human cells and tissues barely contain free CI, which is rather contained in SC I_1_+III_2_ and the respirasome [29, 37]. In contrast, other mammalian mitochondria (from bovine, ovine, rat or mouse) contain larger amounts of CI in its free form, even if the majority is still mostly in the form of SCs [2, 17, 28, 43–45]. The distribution of the MRC complexes between free complexes and SCs seems to differ even more in non-mammalian animal species. Several reptile species contain a very stable SC I+III_2_ that lacks CIV [28], and in *Drosophila melanogaster* practically all of CI is free, SCs being almost completely absent [46, 47]. However, comparative studies of MRC function in diverse animal species suggest that higher amounts and stability of the SCs do not correlate with increased respiratory activity/efficiency and/or reduced ROS production [28, 47].

Here we show that SCs can be stably formed in *D. melanogaster* mitochondria upon mild perturbations of individual CIV, CIII_2_ and CI biogenesis. This finding enabled us to test whether increased SC formation translated into enhanced respiration proficiency. However, MRC performance of fruit fly mitochondria did not change regardless of the presence or absence of SCs. These observations have led us to conclude that: 1) the efficiency in the assembly of the individual complexes is likely to be the main determinant of SC formation and 2) these supramolecular complexes play a more relevant role in maintaining the stability and/or supporting the biogenesis of the MRC than in promoting catalysis.

## RESULTS

### *D. melanogaster* MRC organization does rely on SC formation under physiological conditions

To obtain a detailed characterization of MRC organization in *D. melanogaster,* we isolated mitochondria from wild-type adults and, after solubilization with digitonin, we performed Blue-Native Gel Electrophoresis (BNGE) followed by mass spectrometry analysis of the gel lanes, using ‘Complexome Profiling (CP)’ [48] and thus obtaining a profile of peptide intensities from most OXPHOS subunits along the electrophoresis lane (Figure 1A). This allowed us to unequivocally determine the identity of the main bands that can be visualized by Coomassie staining of BNGE gels (Figure 1B). The identity of the bands corresponding to complex I (CI) and complex IV (CIV) was also confirmed using specific in-gel activity (IGA) assays (Figure 1B). These analyses verified that *D. melanogaster* mitochondria contain extremely low amounts of high-molecular weight CI-containing SCs [46, 47], using the same solubilization and electrophoresis conditions in which the SCs are readily detectable in mammalian cells and tissues [29, 43, 49]. As previously described [47], dimeric complex V (CV_2_) is easily visualized by BNGE and present in similar amounts as monomeric CV in *D. melanogaster* mitochondrial membranes solubilized with digitonin (Figure 1A, B). This CV_2_ species is not a strongly bound dimer, as it disappears when the samples are solubilized using a harsher detergent such as dodecylmaltoside (DDM) [47] (Figure 1C). Conversely, CI is mainly found as a free complex in the native gels irrespective of whether the mitochondria had been solubilized with digitonin or DDM (Figure 1A, B and C). The other minor CI-containing band corresponding to the fraction associated with CIII_2_, accounts for about 3% of the total amounts of CI and CIII_2_, according to label-free mass spectrometry quantifications in the CP analysis and densitometric quantifications of the CI-IGA signals (Figure 1A and Figure S1A). CIV activity is absent in this band both in digitonin- and DDM solubilized samples (Figure 1B, C), whereas it is present in the bands that correspond to individual CIV and dimeric CIV (CIV_2_) detected both in digitonin- and DDM-treated samples, as well as in SC III_2_IV_1,_ which is present only in the digitonin-solubilized samples. This is different from mammalian mitochondria in which SC III_2_IV_1_ is present also in DDM-solubilized mitochondria, probably due to the tight binding of CIII_2_ to CIV through COX7A2L/SCAF1 [4, 29, 50]. The latter does not have a homolog in *D. melanogaster* even though this species has three different COX7A isoforms (named COX7A, COX7AL and COX7AL2) that exhibit a tissue-specific expression pattern, according to FlyBase (www.flybase.org). CP analysis of *Drosophila* mitochondria only detected COX7A (mammalian COX7A1 homolog) and COX7AL2 (mammalian COX7A2 homolog), whereas COX7AL, that is solely expressed in testis, was not found. Therefore, SC I_1_III_2_ can be considered the only stable SC species in physiological conditions in *D. melanogaster*, yet containing a minute fraction of the total CI and CIII_2_.

**Figure 1.**
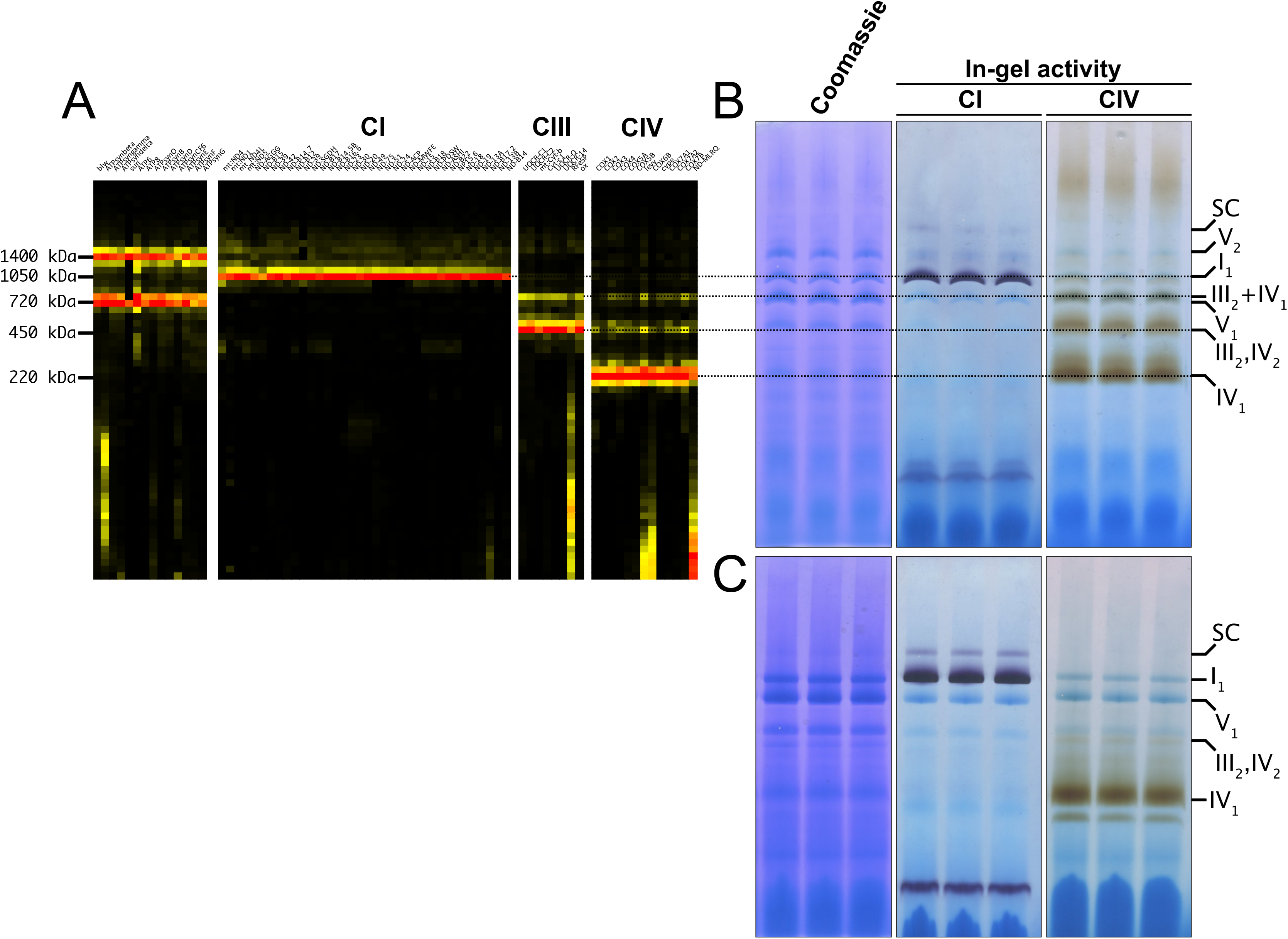
*D. melanogaster* mitochondrial respiratory chain does not rely on SC formation under physiological conditions. **(A)** Complexome profiling of wild-type *D. melanogaster* mitochondria. Heatmaps show relative abundance of MRC subunits belonging to complex I (CI), complex II (CII), complex III_2_ (CIII) and complex IV (CIV). Color scale of normalized peptide intensities are 0 (black), 96^th^ percentile (yellow) and 1 (red). **(B)** BN-PAGE separation of mitochondria from wild-type *D. melanogaster* solubilized with digitonin. Native gels were either stained with Coomassie R250 or analyzed by in-gel activity (IGA) for complex I (CI) and complex IV (CIV). **(C)** BN-PAGE separation of mitochondria from wild-type *D. melanogaster* solubilized with dodecylmaltoside (DDM). Native gels were either stained with Coomassie R250 or analyzed by in-gel activity assay (IGA) for complex I (CI) and complex IV (CIV).

### Perturbations of CIV assembly result in increased formation of SC I_1_III_2_

COA8 is a CIV assembly factor the defects of which cause isolated mitochondrial CIV deficiency in human and mouse [51, 52], as well as in *Drosophila melanogaster* [53, 54]. Consistent with the role of Coa8 in CIV biogenesis, CP analysis of mitochondria from *Coa8* knockout (*Coa8^KO^*) flies showed a clear decrease in fully assembled CIV and in all the CIV containing species (Figure 2A, B) when compared to the corresponding wild-type (WT) individuals (Figure 1A and Figure 2B). Curiously, CP also showed that in the *Coa8^KO^* mitochondria the amounts of SC I_1_III_2_ were noticeably increased (Figure 2A, B). In these samples, complexes I and III_2_ build a stable SC species containing ∼16% of the total amount of CI, as visualized by CI-IGA, as well as by western blot (WB) and immunodetection of specific CI and CIII_2_ subunits after BNGE in DDM-solubilized mitochondria (Figure 2C, D, Figure S1A and S2A). We initially speculated that the ∼5-fold increase of SC I_1_III_2_ formation could be linked to the release of III_2_ from SC III_2_IV_1_ induced by the strong reduction in CIV amounts when Coa8 is absent.

**Figure 2.**
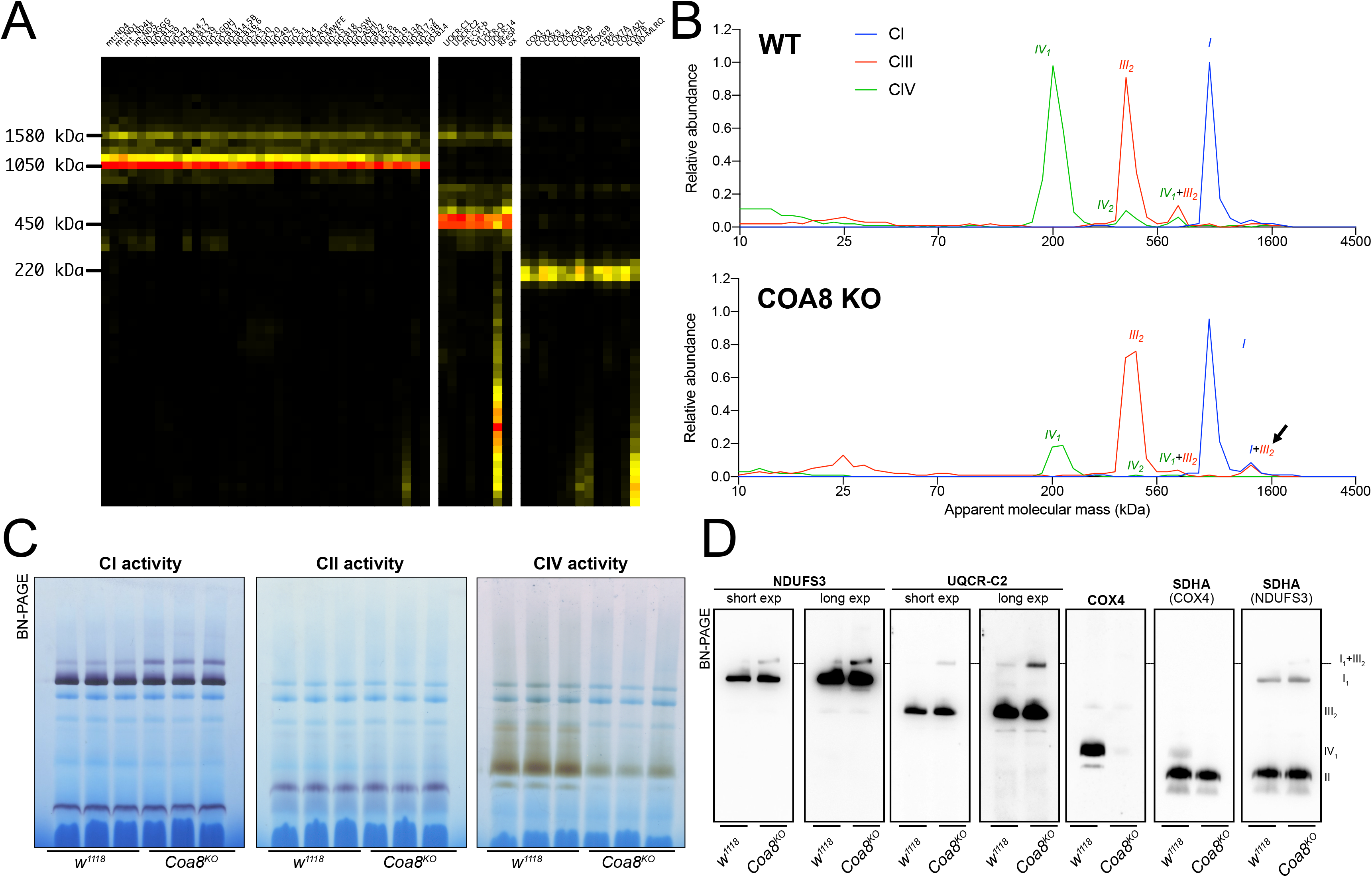
Severely perturbed CIV assembly results in increased formation of SC I_1_III_2_. **(A)** Complexome profiling of Coa8 KO *D. melanogaster* mitochondria. Heatmaps show relative abundance of MRC subunits belonging to complex I (CI), complex II (CII), complex III_2_ (CIII) and complex IV (CIV). Color scale of normalized peptide intensities are 0 (black), 96° percentile (yellow) and 1 (red). **(B)** Average MS profiles depicted as relative abundance of MRC enzymes in natively separated complexes from wild-type (top) and Coa8 KO (bottom) fly mitochondria. Profiles of complexes I, III_2_ and V (CI, CIII and CV) are plotted as average peptide intensity of the specific subunits identified by MS for each complex vs. apparent molecular weight. The increase in the relative abundance of SC I_1_III_2_ in Coa8 KO mitochondria is indicated by a black arrow. **(C)** In gel-activity assays for MRC complex I (CI), complex II (CII) and complex IV (CIV) in DDM-solubilized mitochondria from wild-type (*w^1118^*) and Coa8 KO (*Coa8^KO^*) flies. **(D)** BN-PAGE, western blot immunodetection of MRC complexes from a pool of three control wild-type (*w^1118^*) and and three Coa8 KO (*Coa8^KO^*) fly mitochondria preparations, using antibodies against specific subunits: anti-UQCRC2 (complex III), anti-NDUFS3 (complex I), anti-COX4 (complex IV) and anti-SDHA (complex II).

To test this hypothesis, we modulated Coa8 expression via UAS/GAL4 system using RNAi driven by a ‘mild’ ubiquitous GAL4 driver (*da-gal4*). With this system, the *Coa8* mRNA levels were reduced to ∼60% of the control (Figure 3A). However, these flies showed comparable levels of fully assembled CIV (Figure 3B, Figure S2B). Interestingly, in this case there was also an increased formation of SC I_1_III_2_ from ∼3% in the control to ∼10% in the mild *Coa8^RNAi^* (Figure 3B, C, Figure S1B). Therefore, both strong and weak perturbations of CIV assembly produce an increased formation of CI-containing SCs in *D. melanogaster,* irrespective of whether they result in CIV deficiency or not.

**Figure 3.**
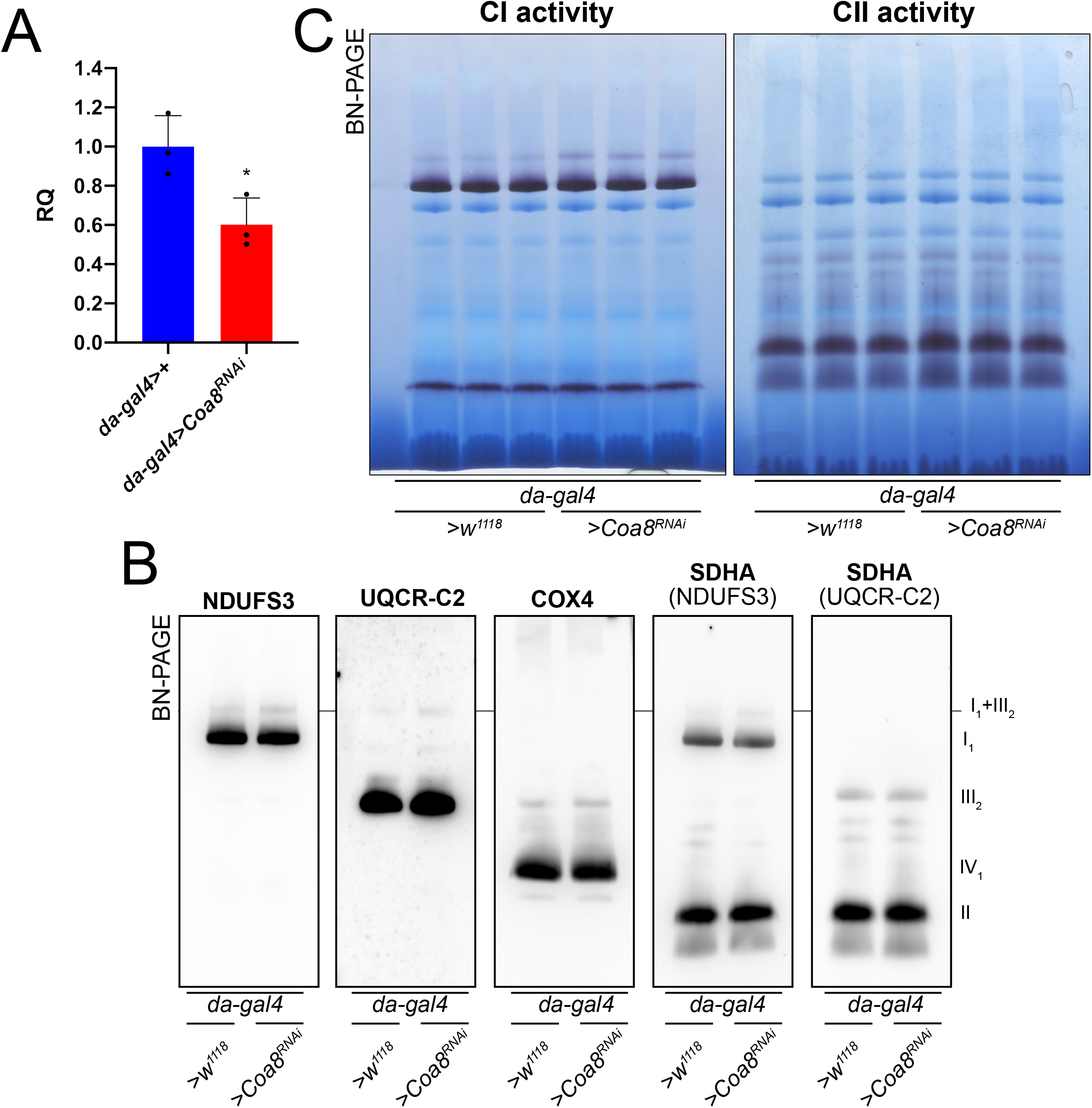
Mildly perturbed CIV assembly results in increased formation of SC I_1_III_2_. **(A)** Relative quantification (RQ) of *Coa8* mRNA expression in control (*da-gal4>+*) and *Coa8* KD (*da-gal4>Coa8^RNAi^*) flies measured by qPCR. Data are plotted as mean[±[SD (n[=[3 biological replicates, Student’s t test *p[≤[0.05). **(B)** In gel-activity assays for MRC complex I (CI), complex II (CII) and complex IV (CIV) in DDM-solubilized mitochondria from control (*da-gal4>+*) and *Coa8* KD (*da-gal4>Coa8^RNAi^*) flies. **(C)** BN-PAGE, western blot immunodetection of MRC complexes from a pool of three control (*da-gal4>+*) and three Coa8 KD (*da-gal4>Coa8^RNAi^*) fly mitochondria samples, using antibodies against specific subunits: anti-UQCRC2 (complex III), anti-NDUFS3 (complex I), anti COX4 (complex IV) and anti-SDHA (complex II).

### Enhanced formation of SC I_1_III_2_ does not result in increased respiratory rates

SC formation was proposed to serve as a means to favor electron transfer between the complexes and therefore increase the efficiency of CI-fueled respiration [1]. With this in mind, the complete *Coa8^KO^* and mild *Coa8^RNAi^* fly mitochondria, which show increased amounts of SC I_1_III_2_ compared to the WT controls, provide an excellent opportunity to test this possibility. Oxygen consumption activities of fly homogenates in the presence of different substrates and inhibitors were analyzed by high-resolution respirometry (Figure 4A, B). The significant decrease in CIV enzymatic activity in the *Coa8^KO^*(Figure 4C) was not reflected by reduced oxygen consumption (Figure 4A). This could be explained as a result of a high CIV excess in fly mitochondria, in which the observed 60% reduction in CIV enzyme activity is still above the threshold at which the CIV defect determines lower respiratory rates [55, 56]. In contrast, the mild reduction in *Coa8* mRNA levels did not result in CIV enzymatic deficiency but produced a slight elevation in CI activity (Figure 4D), which is most likely due to the increased SC I_1_III_2_ formation (Figure 3B and C, Figure S1B). However, the *Coa8-KD* mitochondria did not show any differences in respiration with either CI-linked or CII-linked substrates. Also, the increased and stable interactions between complexes I and III_2_ in the Coa8 deficient models did not produce a preferential utilization of electrons coming from CI, which would be the prediction if SC formation increased electron transfer efficiency [13].

**Figure 4.**
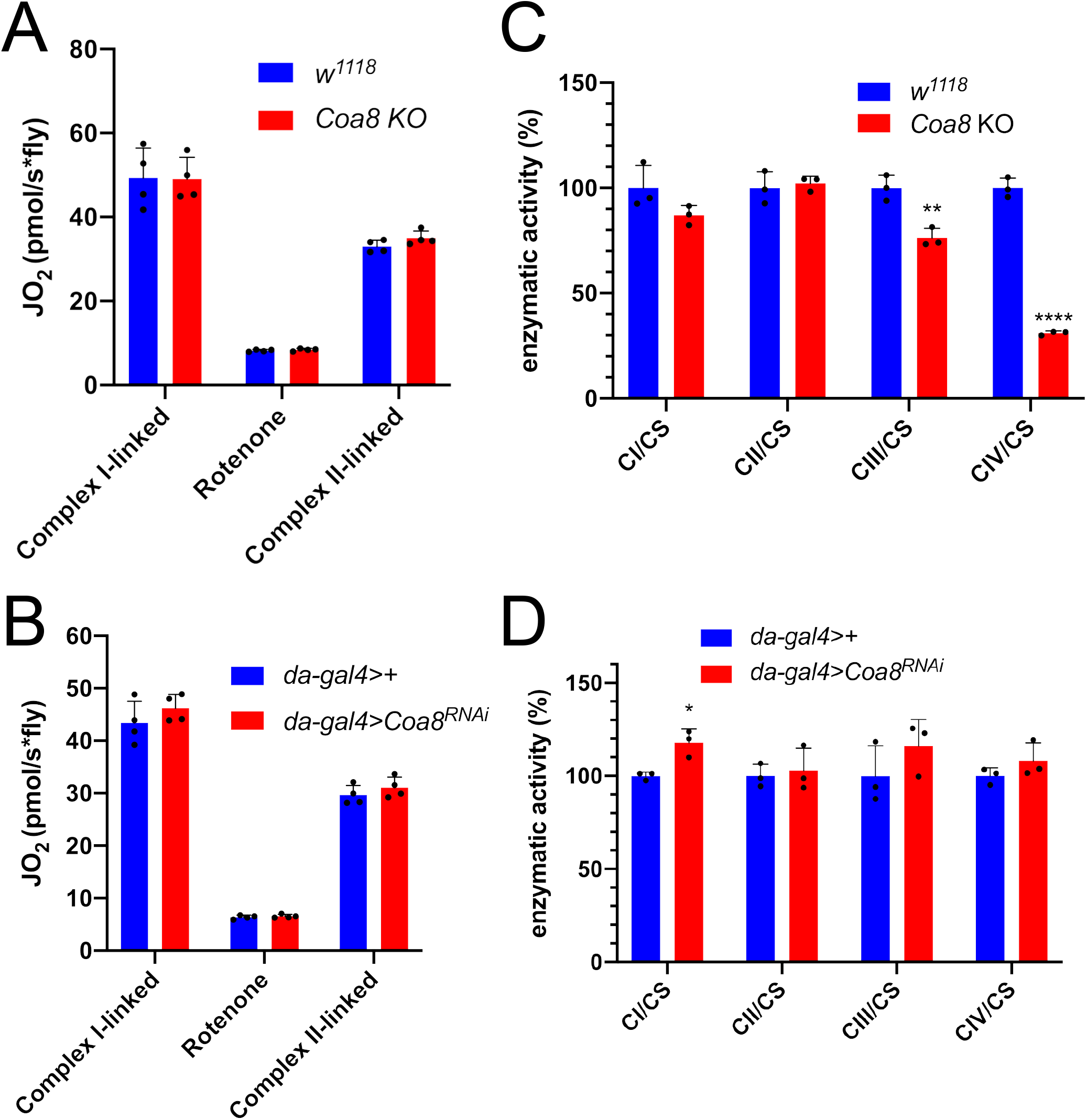
Enhanced formation of SC I_1_III_2_ does not result in increased respiration. **(A-B)** High-resolution respirometry (HRR) analyses of whole-fly homogenates. Respiration is represented by oxygen flux (JO_2_) measured by oxygen consumption rates (OCR – pmol/s*fly). OCR have been measured via substrate-driven respiration under saturating concentrations of substrates inducing either complex I (CI) or complex II (CII) -linked respiration. Rotenone was used to block CI-linked respiration before measuring CII-linked respiration. HRR was performed on **(A)** *Coa8* KO and **(B)** *Coa8* KD fly homogenates compared to relative controls. Data are plotted as mean[±[SD (n[=[4 biological replicates). **(C-D)** Kinetic enzyme activity of individual MRC complexes in **(C)** *Coa8* KO and **(D)** *Coa8* KD compared with the relative control individuals, normalized by citrate synthase (CS) activity. Data are plotted as mean[±[SD (n[=[3 biological replicates, pairwise comparisons by unpaired Student’s t test *p[≤[0.05, **p[≤[0.01, ****p[≤[0.0001).

### Mild perturbation of CIII_2_ biogenesis also enhances SC formation in *D. melanogaster*

To determine whether increased SC I_1_III_2_ formation was specific for CIV deficient flies, we targeted CIII_2_ by knocking down the expression of *Bcs1*. BCS1L, the human homolog, is fundamental for a correct CIII_2_ biogenesis, being responsible for the incorporation of the catalytic subunit UQCRFS1 in the last steps of CIII_2_ maturation [57, 58]. To obtain a severe CIII_2_ defect in *D. melanogaster*, we crossed a ‘strong’ ubiquitous GAL4 driver (*act5c-gal4*) line with a *UAS-Bcs1* RNAi responder line [59]. The knockdown efficiency was high, with a ∼75% decrease in *Bcs1* expression at the mRNA level (Figure 5A). In this model, *D. melanogaster* development was severely impaired causing an arrest at the larval stage [59]. The strong *Bcs1^RNAi^* caused also a significant decrease in fully assembled CIII_2_ levels (Figure 5B, C, Figure S2C) and in CIII_2_ enzymatic activity of about 50% (Figure 5D). Consistent with the observed CIII_2_ deficiency, both the CI- and CII linked respiration rates were significantly decreased by around 40% (Figure 5E).

**Figure 5.**
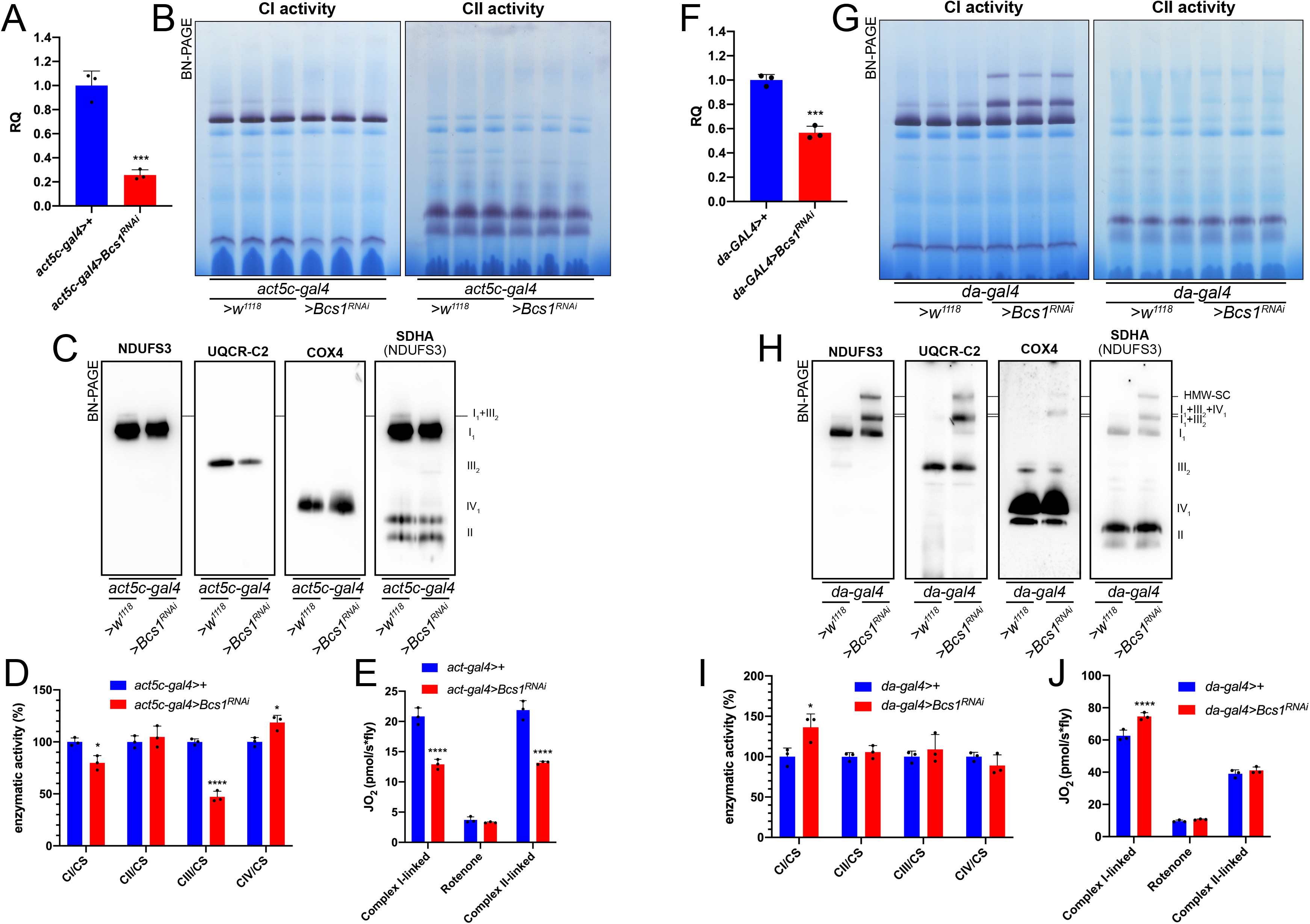
Mild perturbation of CIII_2_ biogenesis enhances SC formation in *D. melanogaster*. **(A)** Relative quantification (RQ) of *Bcs1* mRNA expression in control (*act5c-gal4>+*) and *Bcs1* KD (*act5c-gal4>Bcs1^RNAi^*) larvae measured by qPCR. Data are plotted as mean[±[SD (n[=[3 biological replicates, Student’s t test ***p[≤[0.001). **(B)** In gel-activity assays for MRC complex I (CI), complex II (CII) and complex IV (CIV) in DDM-solubilized mitochondria from control (*act5c-gal4>+*) and *Bcs1* KD (*act5c-gal4>Bcs1^RNAi^*) larvae. **(C)** BN-PAGE, western blot immunodetection of MRC complexes from a pool of three control (*act5c-gal4>+*) and three *Bcs1* KD (*act5c-gal4>Bcs1^RNAi^*) larvae mitochondria samples, using antibodies against specific subunits: anti-UQCRC2 (complex III), anti-NDUFS3 (complex I), anti-COX4 (complex IV) and anti-SDHA (complex II). **(D)** Kinetic enzyme activity of individual MRC complexes in control (*act5c-gal4>+*) and *Bcs1* KD (*act5c-gal4>Bcs1^RNAi^*) larvae mitochondria normalized by citrate synthase (CS) activity. Data are plotted as mean[±[SD (n[=[3 biological replicates, pairwise comparisons by unpaired Student’s t test, *p[≤[0.05, ****p[≤[0.0001). **(E)** High resolution respirometry (HRR) analyses of whole-fly homogenates. Respiration is represented by oxygen flux (JO_2_) measured by oxygen consumption rates (OCR – pmol/s*fly). OCR have been measured via substrate-driven respiration under saturating concentrations of substrates inducing either complex I (CI) or complex II (CII) -linked respiration. Rotenone was used to block CI-linked respiration before measuring CII-linked respiration. HRR was performed on control (*act5c-gal4>+*) and *Bcs1* KD (*act5c-gal4>Bcs1^RNAi^*) homogenates compared to relative controls. Data are plotted as mean[±[SD (n[=[3 biological replicates, two-way ANOVA with Sidak’s multiple comparisons, ****p[≤[0.0001). **(F)** Relative quantification (RQ) of *Bcs1* mRNA expression in control (*da-gal4>+*) and *Bcs1* KD (*da-gal4>Bcs1^RNAi^*) larvae measured by qPCR. Data are plotted as mean[±[SD (n[=[3 biological replicates, Student’s t test ***p[≤[0.001). **(G)** In gel-activity assays for MRC complex I (CI), complex II (CII) and complex IV (CIV) in DDM-solubilized mitochondria from control (*da-gal4>+*) and *Bcs1* KD (*da-gal4>Bcs1^RNAi^*) larvae. **(H)** BN-PAGE, western blot immunodetection of MRC complexes from a pool of three control (*da-gal4>+*) and three *Bcs1* KD (*da-gal4>Bcs1^RNAi^*) larvae mitochondria samples using antibodies against specific subunits: anti-UQCRC2 (complex III), anti NDUFS3 (complex I), anti-COX4 (complex IV) and anti-SDHA (complex II). HWM-SC: high molecular weight supercomplex. **(I)** Kinetic enzyme activity of individual MRC complexes in control (*da-gal4>+*) and *Bcs1* KD (*da-gal4>Bcs1^RNAi^*) larvae mitochondria normalized by citrate synthase (CS) activity. Data are plotted as mean[±[SD (n[=[3 biological replicates, pairwise comparisons by unpaired Student’s t test, *p[≤[0.05). **(J)** High resolution respirometric (HRR) analyses of whole-fly homogenates. Respiration is represented by oxygen flux (JO_2_) measured by oxygen consumption rates (OCR - pmol/s*fly). OCR have been measured via substrate-driven respiration under saturating concentrations of substrates inducing either complex I (CI) or complex II (CII) -linked respiration. Rotenone was used to block CI-linked respiration before measuring CII-linked respiration. HRR was performed on control (*act5c-gal4>+*) and *Bcs1* KD (*act5c-gal4>Bcs1^RNAi^*) homogenates compared to relative controls. Data are plotted as mean[±[SD (n[=[3 biological replicates, two-way ANOVA with Sidak’s multiple comparisons, ****p[≤[0.0001).

In contrast, less pronounced decreases in *Bcs1* expression (Figure 5F) by using the mild *da-gal4* driver instead, did not produce a noticeable CIII_2_ enzymatic defect (Figure 5I). However, the mild *Bcs1-KD* mitochondria showed a very different pattern of CI distribution than the controls (Figure 5G, H, Figure S1D), with the formation of a ‘respirasome-like’ SC I_1_III_2_IV_1_, and also of a higher molecular weight supercomplex (HMW-SC), of unknown stoichiometry, containing also complexes I, III_2_ and IV, as revealed by WB and immunodetection analyses (Figure 5G, H). At the functional level, this was associated with a ∼1.5-fold increase in CI enzymatic activity (Figure 5I), which is proportional to the increase in total CI amounts (Figure S1D), and in higher CI-linked respiration rates but only by ∼1.2-fold (Figure 5J). CII-linked respiration was the same in the mild *Bcs1^RNAi^* samples as in the controls. Therefore, the formation of respirasome-like SCs in these mitochondria did neither increase the efficiency of electron transfer from CI nor determine a diversion of the electronic routes giving preference to the SC-bound CI.

### Mild perturbation of CI biogenesis also leads to increased SC assembly

To understand the effect of the strong and mild perturbations in CI biogenesis on MRC organization in D. melanogaster, we employed a similar strategy as that for CIII (see above). Crossing the strong ubiquitous *act5c-gal4* driver fly line with the *UAS-Ndufs4* RNAi responder line, produced a decrease in *Ndufs4* mRNA expression of ∼90% (Figure 6A). Defects in *NDUFS4* are a major cause of CI deficiency-associated mitochondrial disease in humans [60] and the mouse and *D. melanogaster* animal models display CI deficiency and pathological phenotypes [61, 62]. Accordingly, the strong reduction in Ndufs4 expression observed in our models resulted in developmental arrest and a significant decrease in fully assembled CI levels by ∼40% (Figure 6B, C, Figure S1E) and in a proportional decrease in NADH:CoQ oxidoreductase enzyme activity in the larvae (Figure 6D). BNGE analysis of the strong *Ndufs4^RNAi^ D. melanogaster* mitochondria, revealed the presence of a CI subassembly, containing the core Ndufs3 subunit (Figure 6C, Figure S2E) but lacking NADH-dehydrogenase activity (Figure 6B). This is similar to what is observed in NDUFS4-deficient human and mouse, which accumulate the so-called ∼830 kDa intermediate lacking the N-module and stabilized by the NDUFAF2 assembly factor [63–65]. In contrast, CIV levels and enzyme activity were significantly increased by 1.5-fold in the strong *Ndufs4-KD* (Figure 6C, D, Figure S2E). CI-linked respiration measured in isolated mitochondria from these flies was significantly lower than in the controls, whereas the CII-linked respiration was comparable to the control (Figure 6E). These observations are compatible with the isolated CI defect displayed by the strong *Ndufs4-KD* flies.

**Figure 6.**
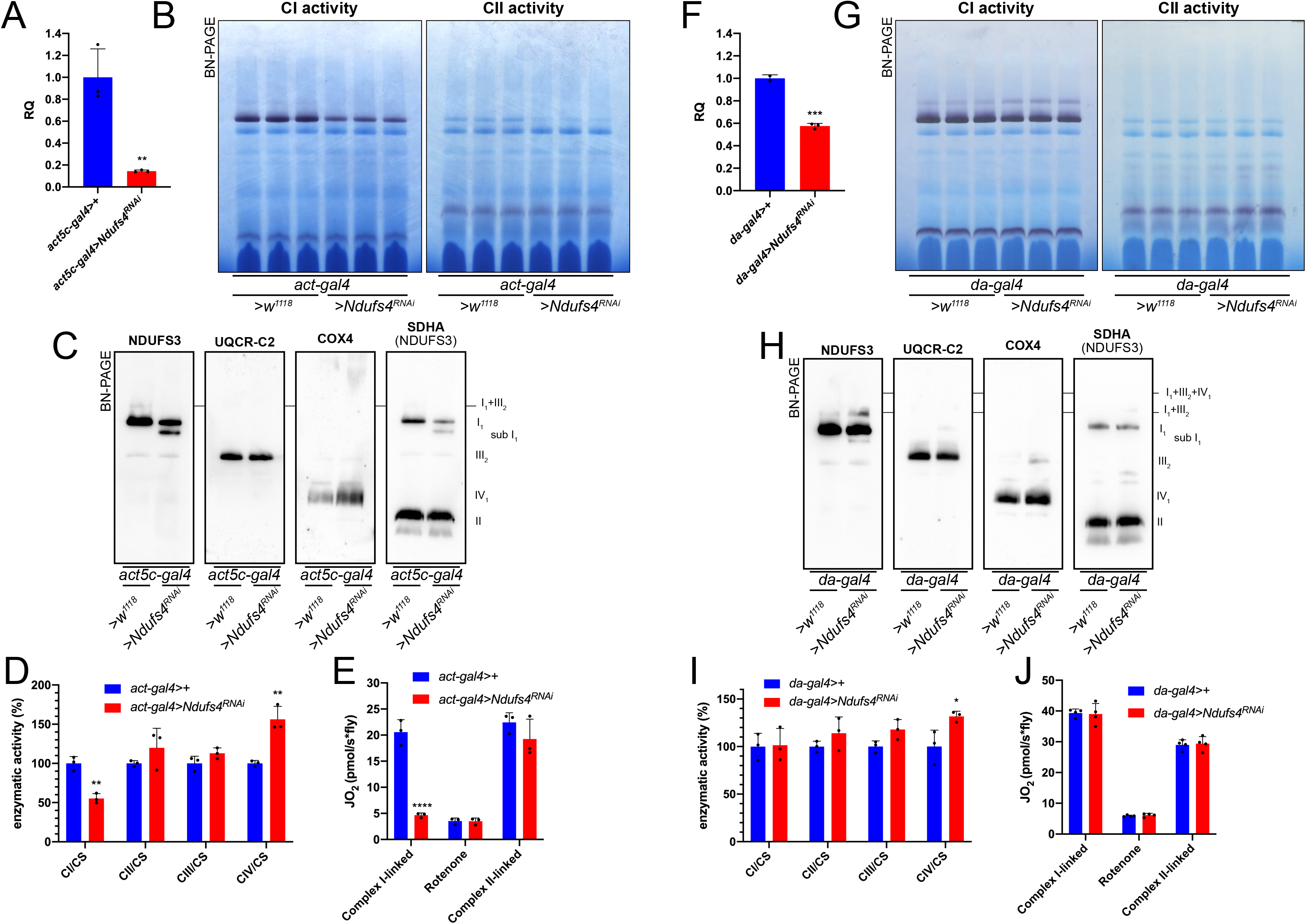
Mild perturbation of CI biogenesis enhances SC formation in *D. melanogaster*. **(A)** Relative quantification (RQ) of *Ndufs4* mRNA expression in control (*act5c-gal4>+*) and *Ndufs4* KD (*act5c-gal4>Ndufs4^RNAi^*) larvae measured by qPCR. Data are plotted as mean[±[SD (n[=[3 biological replicates, Student’s t test **p[≤[0.01). **(B)** In gel-activity assays for MRC complex I (CI), complex II (CII) and complex IV (CIV) in DDM-solubilized mitochondria from control (*act5c-gal4>+*) and *Ndufs4* KD (*act5c-gal4>Ndufs4^RNAi^*) larvae. **(C)** BN-PAGE, western blot immunodetection of MRC complexes from a pool of three control (*act5c-gal4>+*) and three *Ndufs4* KD (*act5c-gal4>Ndufs4^RNAi^*) larvae mitochondria samples, using antibodies against specific subunits: anti-UQCRC2 (complex III), anti-NDUFS3 (complex I), anti-COX4 (complex IV) and anti-SDHA (complex II). **(D)** Kinetic enzyme activity of individual MRC complexes in control (*act5c-gal4>+*) and *Ndufs4* KD (*act5c-gal4>^RNAi^*) larvae mitochondria normalized by citrate synthase (CS) activity. Data are plotted as mean[±[SD (n[=[3 biological replicates, pairwise comparisons by unpaired Student’s t test, **p[≤[0.01). **(E)** High-resolution respirometry (HRR) analyses of whole-fly homogenates. Respiration is represented by oxygen flux (JO_2_) measured by oxygen consumption rates (OCR – pmol/s*fly). OCR have been measured via substrate-driven respiration under saturating concentrations of substrates inducing either complex I (CI) or complex II (CII) -linked respiration. Rotenone was used to block CI-linked respiration before measuring CII-linked respiration. HRR was performed on control (*act5c-gal4>+*) and *Ndufs4* KD (*act5c-gal4>Ndufs4^RNAi^*) homogenates compared to relative controls. Data are plotted as mean[±[SD (n[=[3 biological replicates, two-way ANOVA with Sidak’s multiple comparisons, ****p[≤[0.0001). **(F)** Relative quantification (RQ) of *Ndufs4* mRNA expression in control (*da-gal4>+*) and *Ndufs4* KD (*da-gal4>Ndufs4^RNAi^*) larvae measured by qPCR. Data are plotted as mean[±[SD (n[=[3 biological replicates, Student’s t test ***p[≤[0.001). **(G)** In gel-activity assays for MRC complex I (CI), complex II (CII) and complex IV (CIV) in DDM-solubilized mitochondria from control (*da-gal4>+*) and *Ndufs4* KD (*da-gal4>Ndufs4^RNAi^*) larvae. **(H)** BN-PAGE, western blot immunodetection of MRC complexes from a pool of three control (*da-gal4>+*) and three *Ndufs4* KD (*da-gal4>Ndufs4^RNAi^*) larvae mitochondria samples, using antibodies against specific subunits: anti-UQCRC2 (complex III), anti-NDUFS3 (complex I), anti COX4 (complex IV) and anti-SDHA (complex II). **(I)** Kinetic enzyme activity of individual MRC complexes in control (*da-gal4>+*) and *Ndufs4* KD (*da-gal4>Ndufs4^RNAi^*) larvae mitochondria normalized by citrate synthase (CS) activity. Data are plotted as mean[±[SD (n[=[3 biological replicates, pairwise comparisons by unpaired Student’s t test, *p[≤[0.05). **(J)** High-resolution respirometry (HRR) analyses of whole-fly homogenates. Respiration is represented by oxygen flux (JO_2_) measured by oxygen consumption rates (OCR - pmol/s*fly). OCR have been measured via substrate-driven respiration under saturating concentrations of substrates inducing either complex I (CI) or complex II (CII) -linked respiration. Rotenone was used to block CI-linked respiration before measuring CII-linked respiration. HRR was performed on control (*act5c-gal4>+*) and *Ndufs4* KD (*act5c-gal4>^RNAi^*) homogenates compared to relative controls. Data are plotted as mean[±[SD (n[=[4 biological replicates, two-way ANOVA with Sidak’s multiple comparisons).

Conversely, when *Ndufs4* mRNA expression was reduced to about half of the control levels (Figure 6F), a milder defect in CI abundance (Figure S1F, Figure S2F), which did not affect either NADH:CoQ oxidoreductase enzymatic activity or respiratory capacity was observed (Figure 6G, H, I, J) in the adult flies. However, smaller amounts of the inactive sub-CI were still detectable (Figure 6H). Interestingly enough, this milder perturbation of CI biogenesis by reducing the amounts of Ndufs4 also produced an increase in the formation of SC I_1_III_2_ (Figure 6G, H), without any changes in respiratory performance compared to the controls (Figure 6J).

## DISCUSSION

The first description of MRC SCs in the early 2000’s led to opposite opinions on whether these were real and functionally relevant entities. On the one hand, researchers argued that the random collision model and diffusion of the individual MRC complexes was well established experimentally and the idea of the SCs did not fit with these observations. On the other hand, others considered that these SCs were real entities and therefore they must have a functional relevance, mainly as a means to enhance electron transfer between the individual complexes. Presently, the existence of the SCs is not debated anymore, especially after the determination of the high resolution structures by cryo electron microscopy (EM), first of the mammalian respirasomes (reviewed in [66]), followed by that of other mammalian SCs [3, 4], and of mitochondrial respiratory chain SCs from other eukaryotic species [39–42, 67, 68]. However, whether SCs provide any catalytical advantage to the MRC or not, is still being debated and opposing views continue to exist [18, 69–72]. The recent resolution of MRC SCs from different eukaryotic species has revealed that the relative arrangement of the complexes within the SCs and the bridging subunits varies substantially depending on the species. Therefore, given the conservation of the structures of the individual complexes, one could argue that if SC formation was of capital importance for MRC function, the way the complexes interact should be strictly conserved as well. Importantly, neither in the mammalian respirasomes, nor in the mammalian and plant SC I_1_III_2_ there is any evidence of substrate channeling, as the CoQ binding sites in CI and CIII_2_ are far apart and exposed to the milieu, in principle allowing the free exchange of CoQ [3, 4, 18, 41]. This is in agreement with different sets of functional data indicating that CoQ is interchangeable between the CI-containing SCs and the rest of the MRC [24, 25, 29, 37]. Therefore, this contrasts with the possible segmentation of the CoQ pool - one dedicated to the respirasome and the other to the FADH_2_-linked enzymes – which has been proposed to take place as a consequence of respirasome formation [12–14]. A further element against the notion that SC formation is essential for mitochondrial function, is the fact that in normal conditions the MRC of *D. melanogaster* is predominantly organized based on individual complexes, as shown by us in this work and by others [46, 47]. The *Drosophila* organization was justified by a tighter packing within the mitochondrial cristae and a higher concentration of the mobile electron transporters, as a way to compensate for the lack of SCs [47]. However, inter-species differences in MRC organization do not appear to be of great relevance to determine the level of functionality [28]. For example, disaggregation of CI from the SCs in *A. thaliana*, where there are normally present, did not affect plant viability [73]. Altogether, these observations argue against the strict requirement of SC formation to maintain MRC function.

The second main proposed role for SC is that of stabilizing and/or favoring the assembly of the individual complexes, especially that of CI [37, 38, 74]. Very recently, the *D. melanogaster* CI structure has been solved [75, 76]. This has provided a structural explanation as to why this complex is majorly found as an individual entity, which seems to be due the fact that the N-terminal domain of the NDUFB4 subunit, present in mammals and responsible for the interaction with CIII_2_ within the SCs, is absent in the in the *D. melanogaster* subunit [76]. Additionally, fruit fly subunit NDUFA11 has an extended C-terminus that results in a tighter binding of the subunit to the rest of the CI membrane arm. NDUFA11 stabilizes the transmembrane helix anchoring the lateral helix of subunit MT-ND5, bridging the two parts of the membrane arm of CI in plants and mammals, SC formation seem to be important for stabilizing the interaction of NDUFA11 in the CI membrane domain instead [3, 5, 41, 76, 77]. However, in this work we show that there are ways to induce the formation of CI-containing SCs in fruit fly mitochondria where they normally do not exist. For example, whereas a strong impairment of CIII_2_ biogenesis resulted in CIII_2_ deficiency, decreased respiration and equal amounts of assembled CI, a mild perturbation of CIII_2_ biogenesis did not result in any MRC deficiency but rather in increased CI total amounts, mainly due to enhanced SC formation. Similarly, strong decreases in Ndufs4 expression resulted in low CI amounts and activity, but milder decreases did not affect CI function although CI was redistributed into supramolecular species. In contrast, both the complete KO and mild KD of *Coa8*, induced a significantly increased formation of CI-containing SCs. Even though the *Coa8^KO^* flies display a significant reduction in CIV amounts and activity, the loss of COA8 is associated with milder phenotypes in humans, mouse and flies compared with the lack or dysfunction of other CIV assembly factors [51, 52, 54]. Therefore, we propose that a partial loss of CIV assembly also induces the formation of SCs in *D. melanogaster*. These observations are in line with the idea that in situations of suboptimal MRC complex biogenesis, formation of CI-containing SCs could be a way to structurally stabilize the system and preserve its function [65, 78]. In the context of the cooperative assembly model, where partially assembled complexes get together before completion forming SC precursors [38], one could envision that slower assembly kinetics would increase the chance of interactions at early stages, letting SC assembly occur in a stable way. Difference in the kinetics of CIV assembly have been observed between human and mouse fibroblasts, which is slower in the human cells [79]. Therefore, differences in assembly kinetics could also explain the observed different amounts and stoichiometries of the MRC SCs in different organisms. This is an interesting open question that will deserve further future investigation. In any case, this enhanced SC formation did not translate in an increase in respiratory function nor in a change in substrate preference in any of the tested models. If CI-containing SC formation enhanced respiratory activity significantly, we should have had detected a noticeable increase in CI-linked respiration in all the models of mild perturbation of CIV, CIII_2_ and CI. Although we did observe higher respiration with CI-linked substrates in the mild *Bcs1*-KD, this was even lower than the increase in CI enzymatic activity and total abundance.

Therefore, we conclude that the main role of SC formation is to provide structural stability to the MRC, principally for CI, rather than to enhance electron transfer between the complexes during respiration.

## KEY RESOURCES TABLE

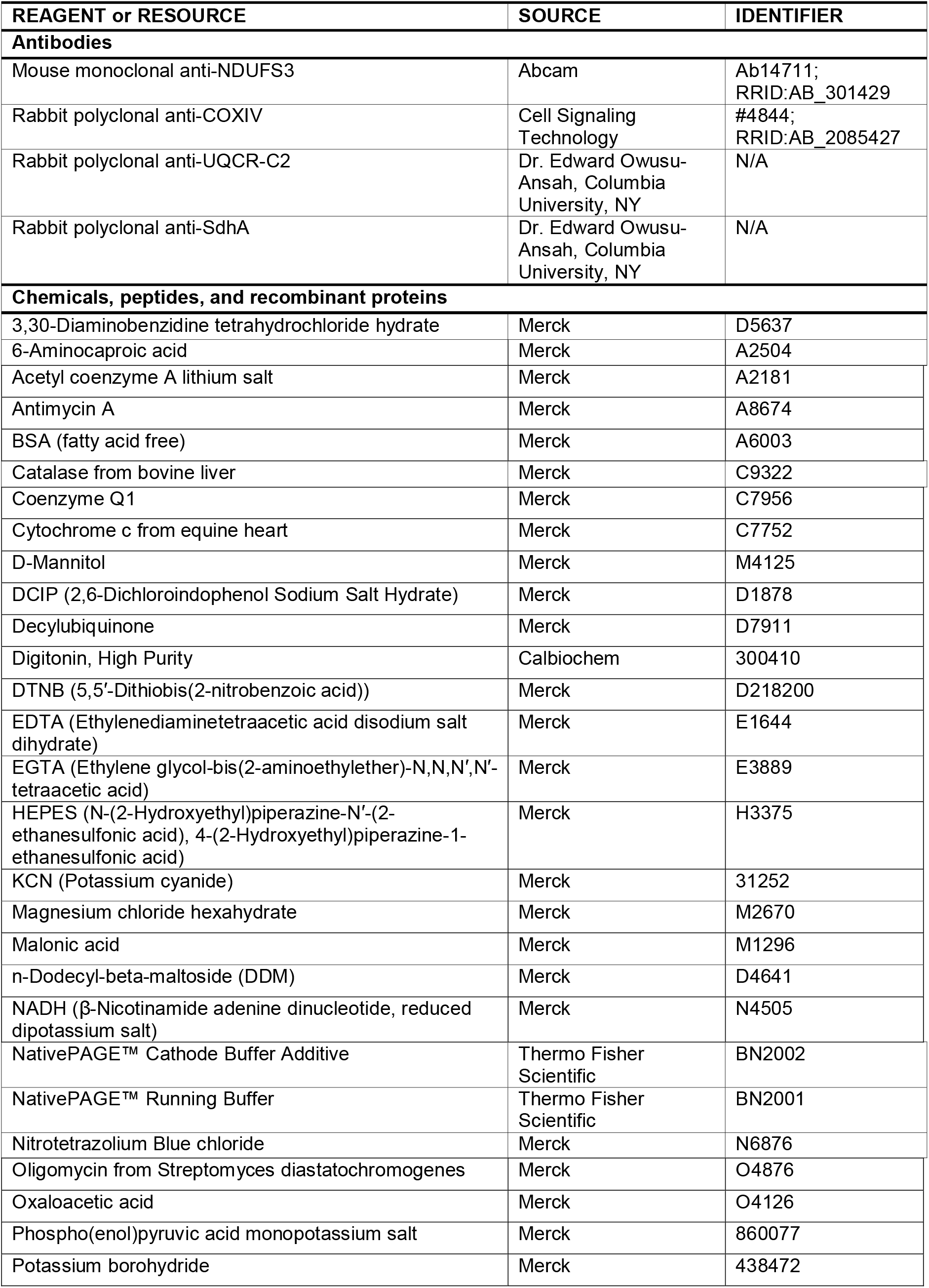
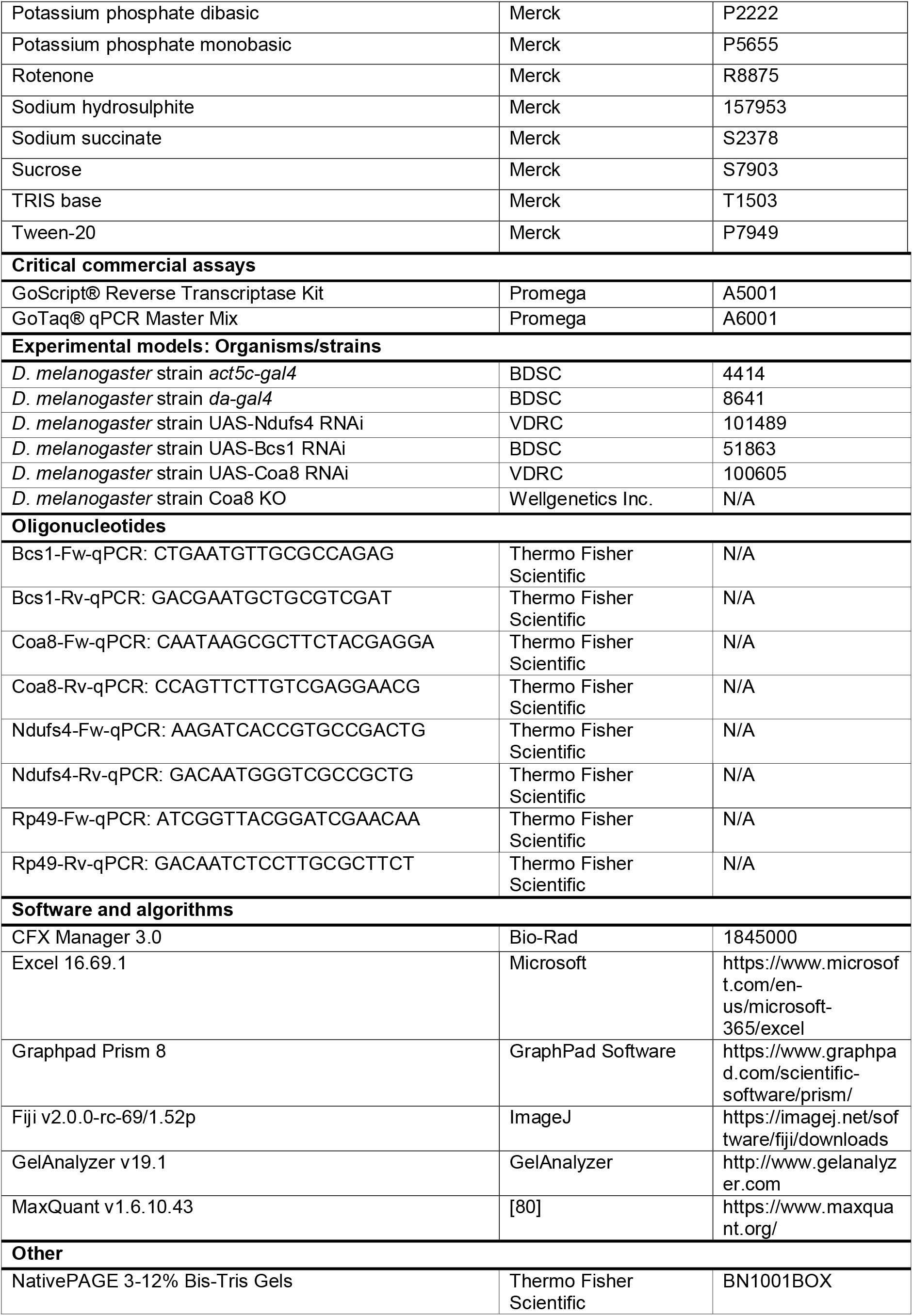
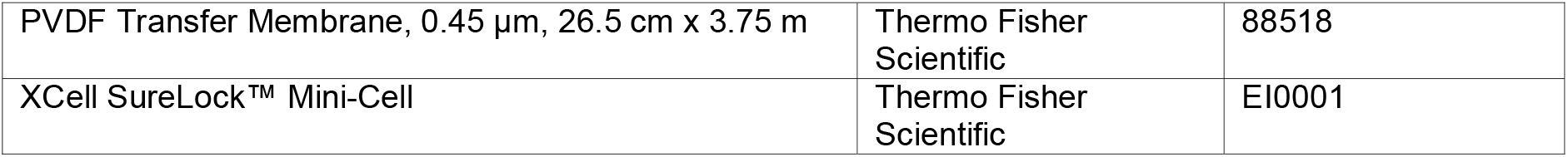

## METHOD DETAILS

### Fly stocks and maintenance

Fly stocks were raised on standard cornmeal medium and kept at 23°C, 70% humidity on a 12:12 hours light/dark cycle. Strains used in this study were obtained from Bloomington Drosophila Stock Center (BDSC) and Vienna Drosophila Resource Center (VDRC). Genotypes used in this study were: *act5c-gal4* (BDSC 4414), *da-gal4* (BDSC 8641), *UAS-Ndufs4 RNAi* (VDRC 101489), *UAS-Bcs1 RNAi* (BDSC 51863), *UAS-Coa8 RNAi* (VDRC 100605). Control strains were obtained in each experiment by crossing the specific *gal4* driver line with the genetic background flies *w^1118^*. *Coa8* KO flies were generated by Wellgenetics Inc. by using CRISPR/Cas9 technology, generating a 676bp deletion, from the -49^th^ nucleotide relative to ATG to the -34^th^ nucleotide relative to the stop codon of Coa8.

### RNA isolation, reverse transcription, and qRT-PCR

Total RNA was extracted from 10 individuals for each genotype using TRIzol method (Thermo Fisher Scientific) according to the manufacturer’s protocol. Reverse transcription was performed with the GoScript Reverse Transcriptase kit (Promega). qRT-PCRs were performed using GoTaq qPCR SYBR Green chemistry (Promega) and a Bio-Rad CFX 96 Touch System (Bio-Rad). The 2^-ΔΔ^Ct method was used to calculate the expression levels of the targets (*Bcs1, Ndufs4, Coa8*) using *Rp49* as endogenous control. The oligonucleotides used are listed in the Key Resource Table

### Isolation of Mitochondria

Mitochondria from *D. melanogaster* larvae and adults were prepared by homogenization and differential centrifugation as described in [81]. Protein concentration of mitochondrial extracts was measured with the Bio-Rad protein assay, based on the Bradford method.

### Blue-native polyacrylamide gel electrophoresis (BN-PAGE) and in-gel activity (IGA) assays

Isolated mitochondria were solubilized in 1.5M aminocaproic acid, 50 mM Bis-Tris/HCl pH 7.0. The samples were solubilized with 4 mg of digitonin (Calbiochem) or 4 mg n-dodecyl β-D-maltoside (Sigma) per mg of protein. After 5 min of incubation on ice, samples were centrifuged at 18,000 X *g* at 4 °C for 30 min. The supernatant was collected and resuspended with Sample Buffer (750 mM aminocaproic acid, 50 mM Bis-Tris/HCl pH 7.0, 0.5 mM EDTA and 5% Serva Blue G). Native samples were separated using NativePAGE 3-12% Bis-Tris gels (Thermo Fisher Scientific) according to the manufacturer’s protocol. For Coomassie staining, gels were stained with Coomassie R 250 for 20 minutes and destained/fixed using 20% methanol, 7% acetic acid. For in-gel activity assays, gels were stained with the following solutions: complex II (succinate dehydrogenase): 5 mM Tris–HCl pH 7.4, 0.2 mM phenazine methosulfate (Sigma), 20 mM succinate, and 1 mg/ml nitrotetrazolium blue chloride; Complex IV (cytochrome *c* oxidase): 50 mM potassium phosphate pH 7.4, 1 mg/ml 3′,3′-diaminobenzidine tetrahydrochloride hydrate (Sigma), 24 units/ml catalase from bovine liver (Sigma), 1 mg/ml cytochrome *c* from equine heart (Sigma), and 75 mg/ml sucrose [82].

### Complexome profiling

Mitochondria from *D. melanogaster* were analyzed by complexome profiling [83]. Isolated mitochondria (0.2 mg) were solubilized with 6 g digitonin/g protein in 50 mM NaCl, 5 mM 6 aminohexanoic acid, 1 mM EDTA, 50 mM imidazole/HCl, pH 7.0. After centrifugation at 22,000 X *g* for 20 min at 4°C, the supernatant was supplemented with Coomassie brilliant blue G250 and proteins were separated by 4-16% gradient BN-PAGE. Digitonin solubilized mitochondrial proteins from bovine heart were loaded as molecular mass standards. Gel lanes were cut into 60 slices, transferred to a 96-well filter microtiter plate (Millipore), and destained in 50% (v/v) methanol, 50 mM ammonium bicarbonate. After destaining, in-gel digestion with trypsin was performed. Tryptic peptides were separated by liquid chromatography and analyzed by tandem mass spectrometry (LC-MS/MS) in a Q-Exactive 2.0 Orbitrap Mass Spectrometer (2.8 SP1) equipped with an Easy nLC1000 nano-flow ultra-high-pressure liquid chromatography system (Thermo Fisher Scientific) at the front end. Thermo Scientific Xcalibur 3.1 Software Package was used for data recording. MS RAW data files were analyzed using MaxQuant (version 1.5.0.25). The extracted spectra were matched against the *Drosophila melanogaster* Uniprot Reference Sequence database (release 2020_04). Database searches were done with 20 ppm match tolerances. Trypsin was selected as the protease with two missed cleavages allowed. Dynamic modifications included N-terminal acetylation and oxidation of methionine. Cysteine carbamidomethylation was set as a fixed modification. Keratins, and trypsin were removed from the list. The abundance of each protein was determined by label-free quantification using the composite intensity based absolute quantification (iBAQ) values determined by MaxQuant analysis and was corrected for loading and MS sensitivity variations between samples based on the total iBAQ value for all detected complex V subunits. Gel migration profiles were created for each protein and normalized to the maximum abundance. Profiles of the identified mitochondrial proteins were hierarchically clustered by distance measures based on Pearson correlation coefficient (uncentered) and the average linkage method using the NOVA software package v0.5 [84]. The visualization and analysis of the heatmaps representing the normalized abundance in each gel slice by a three-color code gradient (black/yellow/red) were done using Microsoft Excel 2019 and Graph Pad Prism 8.4.3. The mass calibration for the BN-PAGE gels was performed as previously described [85]. Membrane proteins were calibrated using the well-known molecular masses of respiratory chain complexes and supercomplexes from bovine heart mitochondria. The soluble proteins were, however, calibrated using the following set of *Drosophila* proteins: ATPB (51 kDa), malate dehydrogenase (72 kDa, dimer), citrate synthase (100 kDa, dimer), ETFA/B (122, heterodimer), heat shock protein 60 (410 kDa, heptamer), ALDH7A1 (675 kDa, dodecamer).

### Western blot and immunodetection

BN-PAGE gels were transferred to PVDF membranes in Dunn carbonate buffer (10 mM NaHCO_3_, 3 mM Na_2_CO_3_) applying a constant current of 300 mA at 4°C for 1 hour using a Mini Trans-Blot® Cell (Bio-Rad). For the immunodetection of specific protein targets, blotted PVDF membranes were blocked in 5% skimmed milk in PBS-T (0.1% Tween-20) at room temperature for 1 hour and then incubated overnight with primary antibodies diluted in 3% BSA in PBS-T overnight at 4°C. PVDF membranes were washed three times with PBS-T for 10 minutes, incubated with the secondary HRP conjugated antibody for 1 hour at room temperature and washed three times with PBS-T for 10 minutes. Chemiluminescent signals were recorded using an Alliance Mini HD9 (UVITEC). Antibodies used are listed in the Key Resource Table. The primary antibodies against *D. melanogaster* UQCR-C2 and SdhA were a kind gift of Dr. Edward Owusu-Ansah (Columbia University, NY).

### High-resolution respirometry

To measure oxygen consumption individuals were homogenized on ice in respiration buffer (120 mM sucrose, 50 mM KCl, 20 mM Tris-HCl, 4 mM KH_2_PO_4_, 2 mM MgCl_2_, 1 mM EGTA, 1% fatty acid-free BSA, pH 7.2). Homogenates were loaded in the chamber of an O2k-HRR (High Resolution Respirometer, Oroboros Instruments) Complex I-linked respiration was measured at saturating concentrations of malate (2 mM), glutamate (10 mM), proline (10 mM) and ADP (2.5 mM). Afterwards, complex II-linked respiration was assessed using 10 mM succinate to the reaction after inhibition of complex I with rotenone (1.25 µM).

### Analysis of MRC enzymatic activities

The activities of mitochondrial respiratory chain complexes and citrate synthase (CS) were measured using kinetic spectrophotometric assays as described [81].

### Statistical analysis

Statistical analysis was performed with GraphPad Prism Software, version 8.2.1. Statistical tests and significance are described in the figure captions.

### Data availability

Complexome profiling data will be deposited in the ComplexomE profiling DAta Resource (CEDAR) repository [86]. Data will be available upon manuscript acceptance.

## ACKNOWLEDGEMENTS

We are grateful to Prof. Rodolfo Costa (CNR institute of Neuroscience, Padova, Italy) for providing the Coa8 KO and Bcs1 and Ndufs4 RNAi lines, Dr. Edward Owusu-Ansah (Columbia University, NY) for sharing the antibodies against *D. melanogaster* UQCR-C2 and SdhA and to Prof. Paolo Bernardi (Dept. of Biomedical Sciences, University of Padova) for critically reading the manuscript.

This research was funded by Fondazione Telethon-Cariplo Alliance GJC21014 (to E.F.-V.), Telethon Foundation GGP19007 (to M.Z.) and GGP20013 (to C.V.), AFM-Telethon 23706 (to C.V.), Department of Biomedical Sciences (University of Padova) FERN_FAR22_01 (to E.F.-V.) and SID2022-VISC_BIRD2222_01 (to C.V.), and Associazione Luigi Comini Onlus (MitoFight2, to M.Z. and C.V.).

## AUTHOR CONTRIBUTIONS

Conceptualization, M.B., M.Z. and E.F.-V.; Methodology, M.B., A.C.-O., S.A. and E.F.-V.; Formal analysis, M.B., A.C.-O., S.A. and E.F.-V.; Investigation, M.B. and A.C.-O.; Data Curation, M.B., A.C.-O., S.A. and E.F.-V.; Writing-Original Draft, M.B. and E.F.-V.; Writing-Review & Editing, M.B., A.C.-O., S.A., M.Z. and E.F.-V.; Visualization, M.B. and E.F.-V.; Supervision, S.A., M.Z., C.V. and E.F-V.; Project Administration, M.B., M.Z., C.V. and E.F.-V.; Funding Acquisition, M.Z., C.V. and E.F.-V.

