## Supplementary material for "Structural rather than catalytic role for mitochondrial respiratory chain supercomplexes": Figure S1A

**Brischigliaro *et al.***

**Figure S1**

**Figure S2**

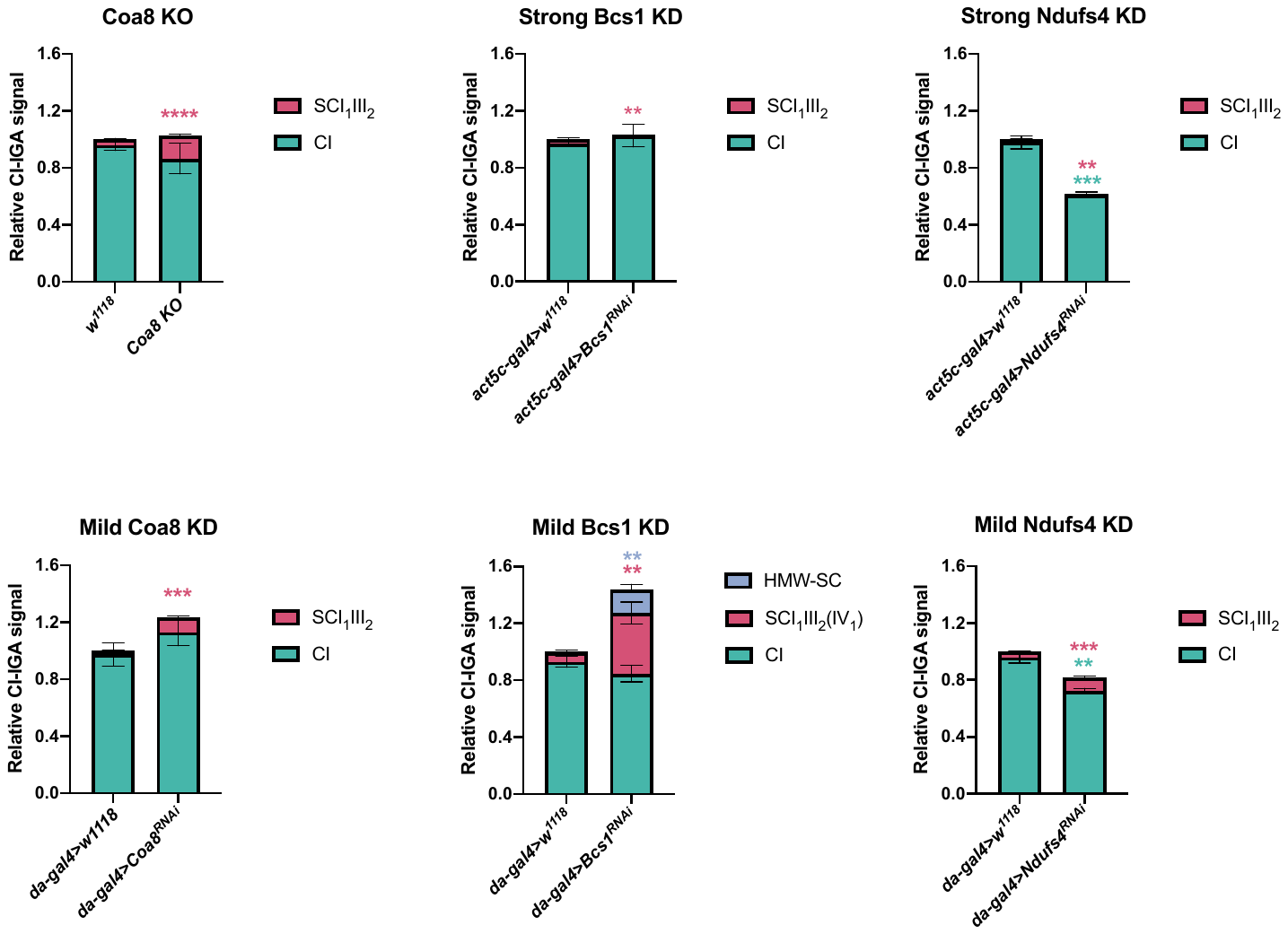

**Supplementary figure 1. Quantification of CI-reactive bands, related to figures 2,3,5 and 6.** The signal intensity of the bands from CI- and CII-in gel activity assays (IGA) of each of the mutated fly strains: (A) *Coa8^KO^*, (B) mild *Coa8^RNAi^*, (C) strong *Bcs1^RNAi^*, (D) mild *Bcs1^RNAi^*, (E) strong *Ndufs4^RNAi^* and (F) mild *Ndufs4^RNAi^* and their corresponding controls were quantified using the Gel analyzer 19.1 software. The graphs show the relative signal intensity of each CI-reactive band normalized to the intensity of the CII-reactive band from the same sample (n=3 biological replicates for each genotype, pairwise comparisons by unpaired Student’s t test **p ≤ 0.01,***p ≤ 0.001, ****p ≤ 0.0001). The total relative intensity of all the CI-containing bands in the control samples was set to 1. Green bars and stars = data corresponding to free complex I; red bars and stars = data corresponding to supercomplex containing CI and CIII_2_ (and CIV in the case of the mild Bcs1 KD); blue bars and stars = data corresponding to supercomplex containing CI, CIII_2_ and CIV. HMW-SC: high molecular weight supercomplex of unknown stoichiometry containing CI, CIII_2_ and CIV.

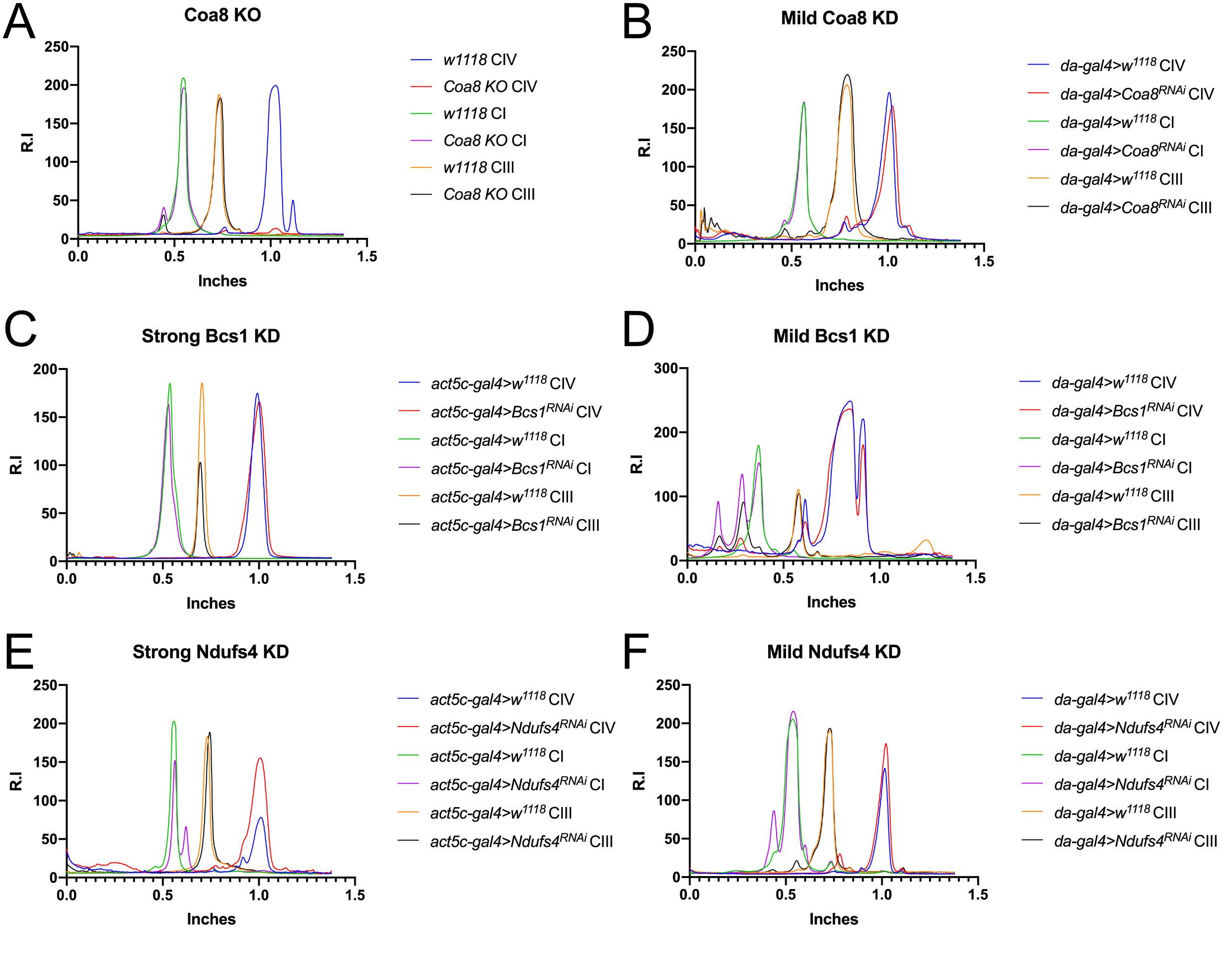

**Supplementary figure 2. Western blot signal profiles for complexes I, III and IV, related to figures 2,3,5 and 6.** WB profiles have been obtained with Fiji (ImageJ) applying the same ROI the BNGE lanes of each genotype: (A) *Coa8^KO^*, (B) mild *Coa8^RNAi^*, (C) strong *Bcs1^RNAi^*, (D) mild *Bcs1^RNAi^*, (E) strong *Ndufs4^RNAi^* and (F) mild *Ndufs4^RNAi^* in comparison with their corresponding controls. Relative intensity (R.I) of the bands along the lane length (in inches) was plotted. High molecular weights are on the left and low molecular weights are on the right.
